# Development of convolutional neural networks for automated brain-wide histopathological analysis in mouse models of synucleinopathies

**DOI:** 10.1101/2025.07.08.663490

**Authors:** A. Barber-Janer, E. Van Acker, E. Vonck, D. Plessers, F. Rosada, C. Van den Haute, V. Baekelandt, W. Peelaerts

## Abstract

Preclinical animal models are indispensable for uncovering disease mechanisms and developing novel therapeutic interventions in synucleinopathies. Key readouts including neuronal cell death, neuroinflammation and alpha-synuclein protein aggregation, are routinely assessed by histological methods. However, traditional characterization of histological samples is labor-intensive and time-consuming. There is a growing need for reproducible and high-throughput tools to capture region- and cell type-specific changes, ultimately improving the predictive value of preclinical studies.

To address this, our study introduces a pipeline using convolutional neural networks (CNNs) for high-throughput, unbiased analysis of immunohistological data in mouse brains. We have trained five CNN-based models to autonomously identify brain regions and detect markers of neurodegeneration, neuroinflammation, and alpha-synuclein aggregation. These models provide accurate, region-specific insights at cellular resolution without manual annotation, significantly speeding up analysis time from weeks to minutes. Our approach enhances the precision and efficiency of histological assessments, providing robust, brain-wide results in various animal models of synucleinopathies.

## Introduction

Neurodegenerative diseases are predicted to become the second leading cause of deaths by 2040^1^. As our population continues to age, substantial efforts are devoted to unraveling the mechanisms that drive neurodegenerative diseases, with the aim of guiding the development of effective therapies. In this context, preclinical models play a central role, and histopathological analysis remains a key approach for assessing disease progression and treatment response at the cellular and tissue level. Unlike clinical neuropathology, which focuses on studying or diagnosing disease in patient brain tissue, preclinical histology aims to systematically characterize disease-relevant features in animal models. Several neuropathological processes, such as neuronal cell death, neuroinflammation, and protein aggregation, are routinely assessed to evaluate therapeutic interventions. However, while essential, their assessment still requires manual counting and annotations, which is labor-intensive and potentially introduces inter-operator variability. As such, conventional immunohistochemical analysis workflows still represent a bottleneck in preclinical research.

In recent years, artificial intelligence (AI) has provided innovative tools in different expert fields allowing fast complex data analysis^2,3^. Deep learning and convolutional neural networks (CNNs) have enabled computers to interpret and analyze image data^4,5^. CNN-based models have been implemented as a novel tool in histopathology to perform automated and unbiased image analysis. These models increase efficiency, improve consistency and enable assessments that are not possible through manual or stereological evaluation. Several CNNs have been developed to detect specific Alzheimer’s disease-related pathological markers, such as amyloid plaque deposition and cerebral amyloid angiopathy in human brain^6,7^. For Parkinson’s Disease (PD), there are recently developed CNN-based models capable of quantifying dopaminergic cells in the substantia nigra^8,9^ or assessing neuroinflammation via the detection and morphological analysis of microglial cells^10^. The results obtained with these models were comparable to manual analysis performed by experienced researchers. Moreover, many of these models can be easily adapted to new data sets. However, all aforementioned models require manual annotation of the regions of interest.

Here we developed a deep learning pipeline using CNNs to perform spatially contextualized immunohistological analysis in mouse brain at cellular resolution. We trained five CNN-based models on brain tissue from mouse models of synucleinopathy. Synucleinopathies are a group of neurodegenerative diseases that include PD and multiple system atrophy (MSA). PD and MSA are characterized by the abnormal accumulation of intracellular inclusions enriched in the protein alpha synuclein (αSyn) in the brains of patients. αSyn protein aggregates and undergoes post-translational modifications such as phosphorylation at Serine 129 (PSer129-αSyn) in patients’ brains^11,12^. In PD, PSer129-αSyn^+^ inclusions are primarily found in neurons as Lewy Bodies, and in neuronal processes as Lewy neurites. MSA patients typically present glial cytoplasmatic inclusions in oligodendrocytes, which are also labelled with the PSer129-αSyn marker. In patients, the presence of αSyn pathology (PSer129-αSyn) correlates with other histopathological observations such as neurodegeneration and neuroinflammation in select brain regions^13,14^.

Our new CNN-based models comprise a set of sequential CNNs, hereafter referred to as layers, designed for the detection of multiple brain regions and neuropathological markers. The first layer, featured in all models, detects brain tissue. The second layer, also present in all models, autonomously identifies distinct brain regions, enabling anatomical localization. Each model then processes this brain region-specific output with a different set of downstream layers, each trained to detect a specific key neuropathological marker, including dopaminergic cells, dopaminergic terminals, neurons, microglia, and αSyn pathology, within the defined regions. We demonstrate that this sequential approach achieves accuracy comparable to conventional stereology while significantly enhancing scalability and reproducibility. We also show that the detection of distinct brain regions can be performed autonomously and reliably. Although in this study the CNN-based models are applied to mouse models of PD and MSA, this methodology is broadly applicable for immunohistochemical analysis of other neuroinflammatory or neurodegenerative *in vivo* mouse models.

## Methods

### Animal procedures

Animal experiments were carried out in accordance with the European Communities Council directive of November 24, 1986 (86/609/EEC) and approved by the Bioethical Committee of KU Leuven (ECD project P008-2021 and P198-2021). 8-week-old female C57BL/6J mice were purchased from Jackson Laboratories.

### Viral vectors and pre-formed fibrils

The recombinant adeno-associated viral (rAAV) vector production and purification was performed by the Leuven Viral Vector Core using the standard triple transfection method as previously described^15^. To produce the neuron-specific vectors, we used a serotype plasmid that coded for the AAV7 capsid and a transgene plasmid containing the wild-type human αSyn cDNA under the control of the CMV enhanced synapsin (CMVeSyn) promoter. To produce the oligodendrocyte-specific vector, we used a serotype plasmid that coded for the AAV-PHP.eB capsid and a transgene plasmid containing the wild-type mouse αSyn cDNA under the control of the myelin-associated glycoprotein (MAG) promoter. The helper plasmid used in both productions was pAdDeltaF6. Real-time PCR was used for genomic copy determination.

Mouse αSyn PFFs were purchased from a commercial vendor (StressMarq Bioscience Inc.). Before injection, PFFs were sonicated with Bioruptor^®^ Pico (BC 100, Diagenode Inc., Belgium), adsorbed onto carbon-coated 200-mesh grids and negatively stained with 2% uranyl acetate. Images of the sonicated PFFs were obtained using transmission electron microscopy and qualitatively assessed to ensure most fragments were shorter than 100 nm. The endotoxin levels were below 0.02 endotoxin units/mg.

### Stereotactic injections

Following anesthesia, 10-week-old C57BL/6J mice were placed in a stereotactic head frame. Injections were performed with a 34-gauge needle and a 10 µl Hamilton syringe at a flow rate of 0.25 μl/min. Stereotactic coordinates were calculated from the skull using bregma as reference. Coordinates to target the right dorsal striatum were: anteroposterior, +0.5 mm; lateral, −2.0 mm; dorsoventral, −3.3 mm. Coordinates to target the right substantia nigra were: anteroposterior: −3.1 mm; lateral, −1.2 mm; dorsoventral, −4.3 mm. All animals were injected with 3µl containing either: PBS for the control groups; 3.9×10^9^ genome copies (GC) of rAAV2/7-CMVeSyn-hαSyn vector in PBS in the right substantia nigra for the neuron-specific overexpression model; 3.9×10^9^ GC of rAAV2/PHP.eB-MAG-mαSyn vector in PBS in the right dorsal striatum for the oligodendrocyte-specific overexpression model; 4 μg of mouse PFFs in PBS the right dorsal striatum for the seeding model.

### Tissue processing and immunohistochemistry

The neuron-specific overexpression model (rAAV2/7-CMVeSyn-hαSyn vector) was sacrificed six months after injection, while the oligodendrocyte-specific overexpression model (rAAV2/PHP.eB-MAG-mαSyn vector) and the seeding model (mouse PFFs) were sacrificed eight months after injection. Age-matched control groups were sacrificed at each timepoint. All C57BL/6J mice were transcardially perfused with ice-cold saline followed by 4% paraformaldehyde (PFA) under non-reversible anesthesia (60mg/kg, Nembutal, Ceva Sante Animale). The full brain was extracted and fixated overnight in 4% PFA. Brains were sectioned at 40 μm thickness in the coronal plane using a vibrating microtome (HM650V, Microm).

Immunohistochemistry (IHC) was performed on free-floating sections. Between all steps three 10 minute washes were performed in 0.1% Tergitol 15-S-9 in PBS (from hereafter referred to as 0.1% Tergitol-PBS). Sections were treated with 3% H_2_O_2_ in PBS for 15 minutes, blocked with 10% goat serum (S26, Sigma-Aldrich) in 0.1% Tergitol-PBS for an hour and then incubated with primary antibodies in 0.1% Tergitol-PBS overnight (listed in **Table 1**). Next day, sections were incubated in biotinylated goat secondary antibody in 0.1% Tergitol-PBS for 2 hours, followed by incubation with Streptavidin-HRP complex (P0397, Agilent Technologies) in 0.1% Tergitol-PBS for 1 hour. Immunoreactivity was visualized using 3,3’-diaminobenzidine (DAB, D5905-50, Sigma-Aldrich) as a chromogen in PBS. Subsequently, sections were dehydrated and mounted. To stain for tyrosine hydroxylase (TH), a marker of dopaminergic cells, the sections were pre-treated with citrate buffer to perform antigen retrieval and before the dehydration procedure substantia nigra sections were counterstained with 0.1% cresyl violet (ab246817, Abcam) to visualize nuclei.

**Table 1.** List of antibodies.

| Antibody | Dilution | Supplier | Catalog number |
| --- | --- | --- | --- |
| Rabbit anti-Tyrosine hydroxylase | 1:10000 | Sigma-Aldrich | AB152 |
| Chicken anti-NeuN | 1:10000 | Sigma-Aldrich | ABN91 |
| Rabbit anti-Iba1 | 1:10000 | Abcam | ab178846 |
| Rabbit anti-pSer129-α-Synuclein | 1:4000 | Abcam | ab51253 (clone EP1536Y) |
| Goat anti-rabbit-biotin | 1:1000 | Abcam | ab6720 |
| Goat anti-chicken-biotin | 1:1000 | Abcam | ab6876 |

### Image acquisition

Whole-slide brightfield imaging was performed with Leica Aperio CS2 scanner (Leica Microsystems) using a 20x objective at 0.22 µm per pixel resolution. Images were acquired in a single plane (no Z-stacks). The digitized images were uploaded to Aiforia® Cloud for training and analysis.

### Training of CNN-based models

Five supervised CNN-based models were trained on the Aiforia® Create platform (AI engine version 7) with whole-slide images containing multiple coronal mouse brain sections. Each section was stained with specific antibodies (listed in **Table 1**) and DAB as the chromogenic substrate to visualize the marker of interest.

All models were built using a layer architecture with multiple nested layers, with each successive child layer analyzing only the pixels provided by the preceding layer in a sequential manner. Aiforia^®^ Create offers two types of layers depending on the nature of the structure to be detected and the hierarchy they hold within a model. Region layers are used to detect areas with a certain type of texture or shape and can have child layers, which means that structures with lower hierarchy can be detected within them. Object layers are used to detect and quantify the number of certain structures and cannot have child layers. Both layer types can be divided into different classes to categorize structures that are at the same level of hierarchy but are independent. For example, brain regions such as “Striatum” and “Motor Cortex” are classes belonging to the “Brain region” layer but are detected independently. Additionally, multiple structures that belong to the same class, hereafter referred to as subregions, are also detected independently within the same slide and are given the name of the class plus a numerical identifier. For instance, one tissue section normally contains two “Striatum” subregions, “Striatum 1” and “Striatum 2”, as this brain region is present in both hemispheres. The additional “Striatum” subregions in all other tissue sections get subsequent identifiers “Striatum 3”, “Striatum 4”, etc. The combination of multiple layers and classes into a single model allows simultaneous detection of targets of interest within various anatomical regions across the whole brain.

A similar annotation strategy was followed to train all five models. **Supplementary Fig. 1** shows a graphic description of the models’ layer hierarchy and annotation examples for each layer. All models feature a common parent layer, “Tissue”, that was annotated to discriminate the brain tissue from the slide background. The training annotations for “Tissue” layer contained: positive tissue, background and regions with edges of tissue and background. In the second layer “Brain region” various brain regions were annotated as separate classes. The output of “Brain region” layer is the value of the area of each subregion detected. An optional child layer under the “Brain region” layer, “Marker^+^ area”, was trained by annotating the positive signal of a specific marker. The readout of this layer is the value of positively stained areas within a specific brain region. This layer is included in the neuronal cell detector (NCD) model and the pSer129-α-synuclein detector (pSynD) model. Lastly, the child layer “Marker^+^ object”, which is under “Brain region” in the dopaminergic cell detector (DCD) model and the microglial cell detector (MCD) and under “Marker^+^ area” in the pSynD model, counts separate objects of interest, such as cells or inclusions. Instance segmentation was used in the DCD model, the MCD model and the pSynD model to obtain each object’s individual morphological metrics. Training annotations for layers “Marker^+^ area” and “Marker^+^ object” included regions with positive signal, containing clusters and isolated objects of various sizes and different morphologies, as well as background signal.

The architecture of each model is described below in **Table 2**. A list of classes included in each layer can be found in **Supplementary Data 1** together with the advanced training and post-training parameters for each model (**Supplementary Data 2-9**).

**Table 2.** List of models and layer architecture.

| | Dopaminergic Cell Detector (DCD) | Dopaminergic Terminal Detector (DTD) | Neuronal Cell Detector (NCD) | Microglia Cell Detector (MCD) | pSer129-α-Synuclein Detector (pSynD) |
| --- | --- | --- | --- | --- | --- |
| Antigen | Tyrosine hydroxylase | Tyrosine hydroxylase | NeuN | Iba1 | pSer129-αSyn |
| Chromogen | DAB | DAB | DAB | DAB | DAB |
| Counterstain | Cresyl violet | None | None | None | None |
| Layer 1 | Tissue (region) | Tissue (region) | Tissue (region) | Tissue (region) | Tissue (region) |
| Layer 2 | Brain region (region) | Brain region (region) | Brain region (region) | Brain region (region) | Brain region (region) |
| Layer 3 | TH<sup>+</sup> cell<sup>s</sup> (object) | - | NeuN<sup>+</sup> cellular area (region) | Iba1<sup>+</sup> cell (object) | pSer129-αSyn<sup>+</sup> area (region) |
| Layer 4 | - | - | NeuN<sup>+</sup> cell (object) | - | Cellular inclusion (object) |
| Segmentation | Yes | No | No | Yes | Yes |

### Validation

All CNN-based models were validated using the Aiforia^®^ Create platform. The performance of each CNN was compared to manual annotations conducted by three experienced validators who were blinded to predefined regions from brain sections which had to be annotated. These regions were not included in the training dataset. The metrics used to evaluate the performance of the models include: precision, calculated as overlap of model analysis results with validator annotations compared to the total model results (TP/(TP+FP)); sensitivity, measured as overlap of model analysis results with the validator annotations compared to the total validator annotations (TP/(TP+FN)); and F1-score, the harmonic mean of precision and sensitivity (2 x precision x sensitivity/(precision + sensitivity))^16^.

The metrics of a validator against each model were averaged and are shown in **Supplementary Fig. 2**. The same metrics were calculated between all validators to evaluate inter-validator variability in **Supplementary Fig. 4**. The F1-score between each pair of validators was then compared to the F1-score of each validator against the model’s results in **Supplementary Fig. 3**. Models were considered ready for use when there were no significant differences between the models’ performance and inter-validator variability. TP (true positive), FP (false positive), FN (false negative).

### Stereological quantification

Unbiased stereology was used to validate the DCD and NCD models. Stereological quantification was performed using the optical fractionator method in a computerized system as previously described (Stereo Investigator; MicroBrightField, MBF)^17^. For the quantification of dopaminergic cells in the substantia nigra, every 5^th^ section throughout the entire SN was analyzed for TH immunoreactivity, leading to seven to eight sections analyzed for each animal. Loss of dopaminergic cells was represented as percentage, comparing the ipsilateral counts to the contralateral counts of each animal. For the quantification of neurons in the dorsal striatum, every 12^th^ section throughout the striatum was analyzed for NeuN immunoreactivity, leading to a total of eight sections analyzed per animal. The coefficients of error, calculated according to the procedure of Schmitz and Hof as estimates of precision, varied between 0.03 and 0.05. All analyses were performed by an investigator blinded to the different conditions.

### Manual dopaminergic terminal immunoreactivity quantification

Between six and seven sections across the whole striatum were stained using an antibody against TH as previously described. Manual annotations of the dorsal striatum and intensity measurements were performed using the software ImageJ. Loss of dopaminergic terminals was represented as percentage, comparing the ipsilateral weighted intensity of the TH staining to the contralateral weighted intensity of each animal.

### Post-processing data output

Data output from each model was post-processed using Python code. For each model a different Python script was developed. These scripts combine data from subregions that belong to the same class and perform model-specific calculations. For the common “Brain region” layer, each subregion detected within multiple tissue sections that corresponds to the same brain region will be merged and processed together. Therefore, the code provides separate results for each brain region for multiple animals simultaneously. Additionally, all Python scripts offer both brain-wide analysis (results from both hemispheres combined) and hemisphere comparison analysis (contralateral vs. ipsilateral) for brain regions that are not physically connected (striatum, substantia nigra, amygdala, globus pallidus, etc.). Select scripts offer the option to include a quality control of the raw data, defined by the researcher. The threshold for removal of a detected object is based on CNN performance confidence and morphological metrics such as area. Exclusion criteria for object removal are listed in **Supplementary Table 1**. The script for each model generates various output readouts to provide a comprehensive overview of the marker assessed (**Supplementary Table 2** and **Supplementary Table 3**).

### Code availability

The underlying Python code for this study is available in GitHub as separate Jupyter Notebooks, which can be accessed via this link: https://github.com/DieterPlessers/Neuro_IHC_analysis.

### Statistical analysis

Statistical analysis was performed using GraphPad Prism 10. The type of test and post hoc correction for multiple testing is indicated in the text or in the legend of each figure. Each dataset was subjected to testing for normal distribution and if not passed, the appropriate non-parametric test was applied.

## Results

Histopathological research relies on key immunohistochemical markers, many of which are routinely applied in preclinical studies of neurodegenerative diseases. We established a high-throughput analysis pipeline that relies on supervised, multi-layered CNNs accessed through a cloud-based platform^10^ to optimize histopathological screenings in preclinical mouse studies of synucleinopathies. We designed a layer-tree architecture for different detection models that allows the autonomous identification of mouse brain tissue on a glass slide and labelling of different brain regions. Subsequently, each model recognizes a specific marker: dopaminergic cells, dopaminergic terminals, neurons, microglia or pSer129-αSyn^+^ inclusions (**Supplementary Fig. 1 & Table 2**).

To demonstrate the effectiveness of our CNN-based models as new tools to screen for neurodegeneration, neuroinflammation and protein inclusion aggregation we employed various mouse models of synucleinopathy (PD and MSA). For these animal models, αSyn pathology is typically induced by selective expression of αSyn protein in either neurons (PD models), or oligodendrocytes (MSA models), via transgenesis or viral vector-mediated transgene expression. Another common method to model PD involves the injection of αSyn pre-formed fibrils (PFFs), which seed further aggregation using the endogenous αSyn present in αSyn-expressing cells. The excessive accumulation of αSyn inclusions results in neurodegeneration and neuroinflammation^18–20^.

### Dopaminergic cell detector

One of the main features of synucleinopathies is the degeneration of the dopaminergic neurons from the nigrostriatal pathway. The loss of dopaminergic cells in the substantia nigra contributes to the development of classical parkinsonian motor symptoms^21,22^. Dopaminergic cell quantification is therefore one of the main readouts used to screen synucleinopathy animal models. We built a CNN-based model to identify dopaminergic cells immunoreactive for tyrosine hydroxylase (TH) (**Fig. 1a**). Images from different PD and MSA mouse models were used, including images from control mice, processed by multiple researchers to introduce variability for model training. Due to the sequential nature of the model’s architecture (**Table 2 & Supplementary Fig. 1a**), we first trained a layer termed “Tissue”, to discriminate brain tissue from the slide’s background, using training examples that included both unannotated background and annotated brain tissue. The second layer was trained to detect the substantia nigra within a brain section. For this training we annotated the substantia nigra within sections that contain this region and left unannotated sections that do not. Lastly, an object layer was built to detect dopaminergic cells. For the training of this layer we annotated dopaminergic cells within the substantia nigra. An object size of 15 μm in diameter and instance segmentation were applied to obtain morphological metrics of each individual cell (**Supplementary Data 3**).

**Figure 1.**
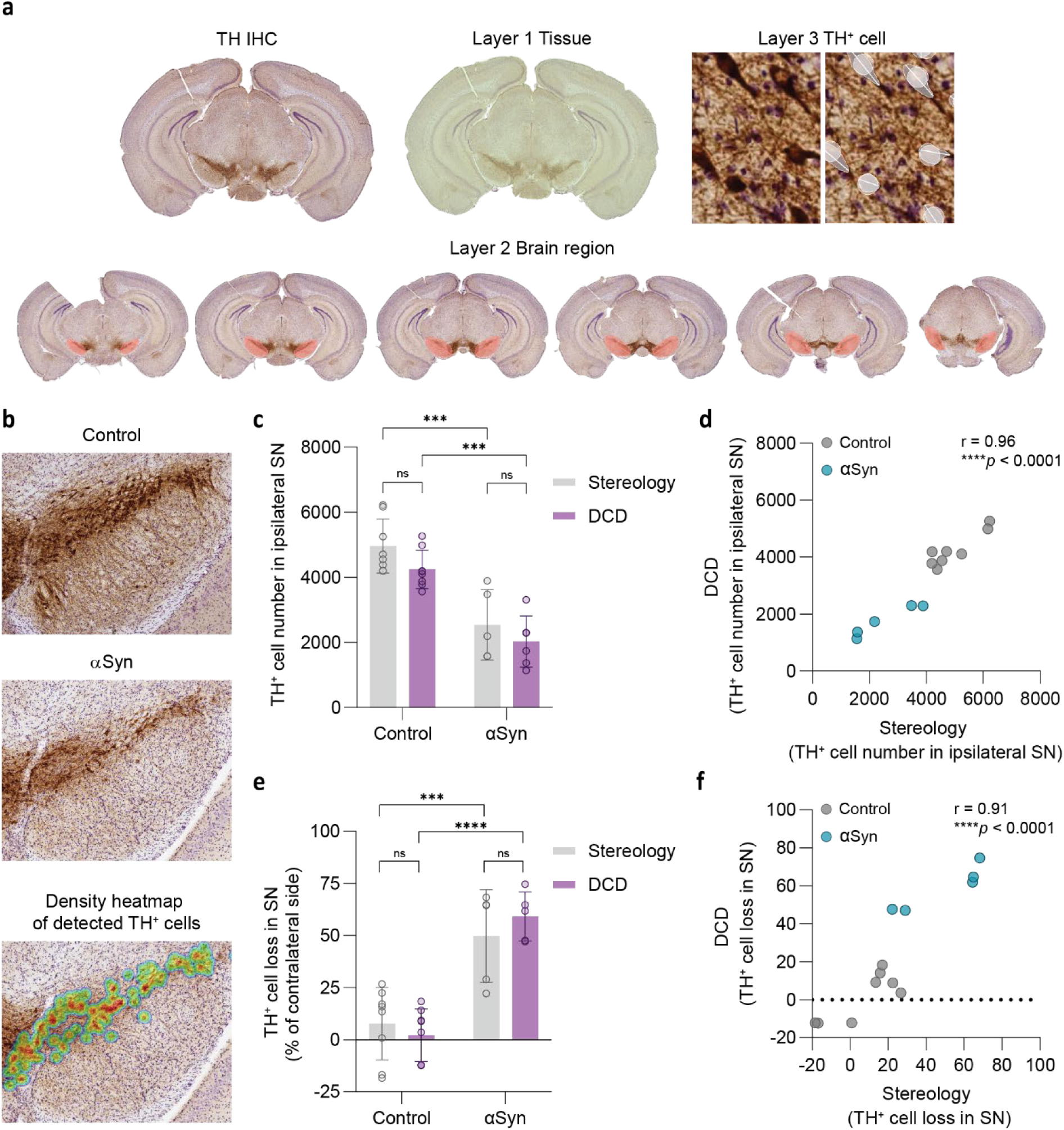
Quantification of dopaminergic cells using the DCD model. Control mice were injected with PBS and αSyn mice were injected with a rAAV vector expressing αSyn in neurons in the substantia nigra. Analysis was performed 6 months post-injection. **a)** Examples of unannotated TH IHC and DCD detection per layer: “Tissue” (layer 1, green), “Brain region” (layer 2, red), and “TH^+^ cell” (layer 3, white). **b)** Representative images of TH and Cresyl Violet staining in the substantia nigra of control and αSyn-overexpressing mice and density heatmap of TH^+^ cells detected by DCD model. **c)** Comparison of TH^+^ cell number in the ipsilateral substantia nigra measured by DCD and stereology (n = 5-8 per group, mean ± SD, two-way ANOVA test with Šídák’s correction for multiple testing, ns, ***p < 0.001). **d)** Correlation between ipsilateral TH^+^ cell number measured by DCD and stereology (Pearson correlation r = 0.96, ****p < 0.0001). **e)** Comparison of TH^+^ cell loss in substantia nigra measured by DCD and stereology (n = 5-8 per group, mean ± SD, two-way ANOVA test with Šídák’s correction for multiple testing, ***p < 0.001, ****p < 0.0001, ns (non-significant)). **f)** Correlation between TH^+^ cell loss measured by DCD and stereology (Pearson correlation r = 0.91, ****p < 0.0001). SD (standard deviation), ns (non-significant).

To evaluate the performance of the dopaminergic cell detector model, hereafter referred to as DCD, we analyzed a set of mice unilaterally injected in the substantia nigra with either phosphate buffered saline (PBS) (referred to as control in **Fig. 1**) or a viral vector expressing αSyn in neurons (referred to as αSyn in **Fig. 1**). This dataset was also analyzed via stereology, the current standard method for cell quantification. We quantified the number of TH^+^ cells in the ipsilateral substantia nigra using both methods and found no statistically significant differences between them, while both methods were able to detect significant differences between the control group and the αSyn-overexpressing group. In control mice, the DCD counted 4,248 ± 598 TH^+^ cells and stereology quantified 4,961 ± 831 TH^+^ cells. In αSyn-overexpressing mice, the DCD found 2026 ± 786 TH^+^ cells and stereology 2541 ± 1083 TH^+^ cells (**Fig. 1c**, two-way ANOVA test with Šídák’s correction for multiple testing, non-significant). Accordingly, when comparing the ipsilateral TH^+^ cell number to the contralateral side for the αSyn group, the DCD model quantified 59.2 ± 11.8 % TH^+^ cell loss in the substantia nigra while stereology measured 49.8 ± 22.2 % TH^+^ cell loss. No neurodegeneration was detected in the control group with either method (DCD: 2.1 ± 12.7 % TH^+^ cell loss; stereology: 7.6 ± 17.3 % TH^+^ cell loss) (**Fig. 1e**, two-way ANOVA test with Šídák’s correction for multiple testing, non-significant). There is significant correlation between both methods in the quantification of TH^+^ cells in the ipsilateral substantia nigra (Pearson’s r = 0.96, \*\*\*\**p* < 0.0001) and TH^+^ cell loss (Pearson’s r = 0.91, \*\*\*\**p* < 0.0001), showcasing the applicability of our new DCD model (**Fig. 1d, f**). Notably, the DCD model provides less variable results than stereology, likely due to its ability to quantify all cells present in a section, bypassing the error-prone sampling used in stereology.

### Dopaminergic terminal detector

During degeneration, the loss of striatal dopaminergic terminals precedes the loss of nigral dopaminergic cell bodies. This degeneration reduces dopamine release and disrupts the basal ganglia’s motor circuits responsible for movement control^21^. We trained a model to detect dopaminergic terminal density in the dorsal striatum of mice (**Fig. 2a**). This serves as a method to assess the integrity of the dopaminergic system and detect early neurodegenerative changes. The dopaminergic terminal detector (DTD) model was trained on coronal brain sections collected from multiple synucleinopathy mouse models (PD and MSA) and control mice, which were processed by different researchers to ensure robust model performance across experimental conditions. The DTD model includes a first layer that detects tissue sections and a second layer that identifies the dorsal striatum (**Table 2 & Supplementary Fig. 1b**). We first trained the “Tissue” layer, using training examples that included both unannotated background and annotated brain tissue. The “Brain region” layer was trained to detect the dorsal striatum within a coronal section. For this training we annotated the striatum within sections that contain this region and left unannotated sections that do not (**Supplementary Data 4**).

**Figure 2.**
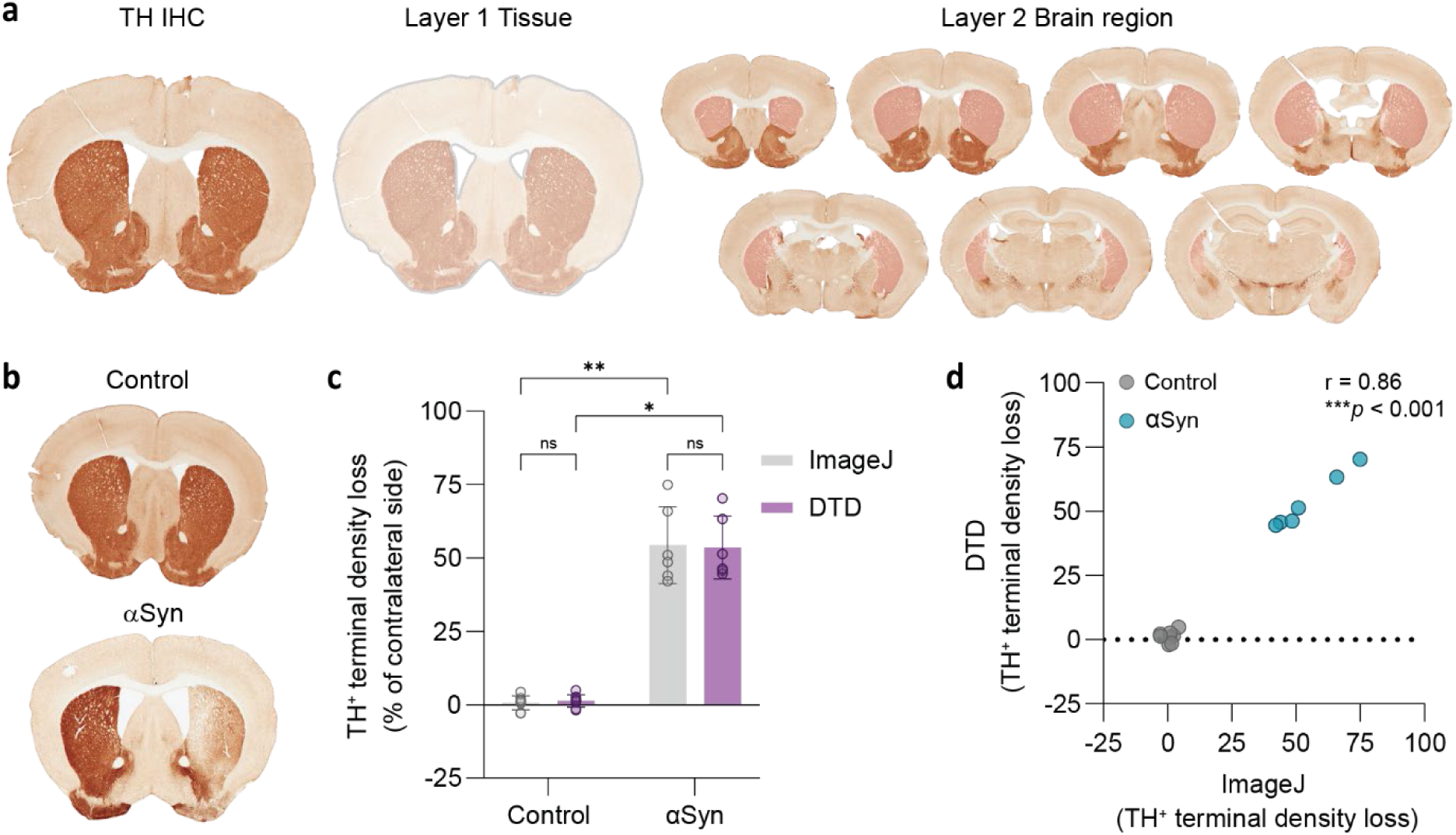
Quantification of dopaminergic terminal density using the DTD model. Control mice were injected with PBS and αSyn mice were injected with a rAAV vector expressing αSyn in neurons in the substantia nigra. Analysis was performed 6 months post-injection. **a)** Examples of unannotated TH IHC and DTD detection per layer: “Tissue” (layer 1, gray) and “Brain region” (layer 2, pink). **b)** Representative image of TH staining in a striatal section from control and αSyn mice. **c)** Comparison of DTD and manual optical density analysis in ImageJ to quantify striatal TH^+^ terminal loss density loss across six sections containing dorsal striatum (n = 6-8 per group, mean ± SD, Kruskal-Wallis test with Dunn’s correction for multiple testing, ns, *p < 0.05, **p < 0.01). **d)** Correlation between striatal TH^+^ terminal density loss measured by DTD and the respective measure obtained with ImageJ (Spearman correlation r = 0.86, ***p < 0.001). SD (standard deviation), ns (non-significant).

To determine the ability of the DTD model to quantify dopaminergic terminal density loss in a unilateral mouse model of PD, we analyzed brain tissue from mice injected with a viral vector expressing αSyn in neurons in the right substantia nigra (referred to as αSyn in **Fig. 2**). The control group was injected with PBS. These are the same groups analyzed previously with the DCD model. We compared the DTD model to manual optical density analysis with ImageJ. The results show that both methods measured comparable loss of TH^+^ terminal density in the ipsilateral dorsal striatum of αSyn mice, with the DTD quantifying a TH^+^ terminal density loss of 53.6 ± 10.7% while ImageJ quantified a 54.4 ± 13.1% decrease (**Fig. 2c**, Kruskal-Wallis test with Dunn’s correction for multiple testing, non-significant). The results from TH^+^ terminal density loss obtained through DTD significantly correlated with results from ImageJ (**Fig. 2d**, Spearman’s r = 0.86, \*\*\**p* < 0.001).

### Neuronal cell detector

Loss of several neuronal types in multiple brain regions is another common feature of PD and MSA^13,23,24^. Assessing the integrity of different brain regions in an animal model can be crucial to link neurodegeneration to behavioral deficits, as well as understanding the relationship between neurodegeneration, neuroinflammation and αSyn inclusion deposition.

To this aim, we developed a neuronal cell detector model, hereafter referred to as NCD, to quantify the number of neurons in whole-slide images using the neuronal marker NeuN. The NCD model was trained to autonomously recognize multiple brain regions in coronal tissue sections and count neuronal cells within these regions (**Table 2 & Fig. 3a**). Brain and spinal cord sections from multiple synucleinopathy mouse models (PD and MSA) and control mice processed by several researchers were included in the training to introduce variability. The “Tissue” layer was annotated as described for previous CNN-based models. The “Brain region” layer was trained with 4621 annotations featuring 28 different brain region classes (**Supplementary Data 1**). Certain brain regions, such as piriform cortex or hippocampus, exhibit such high neuronal density that counting individual cells becomes unfeasible. To address this, we trained a layer to quantify the NeuN positively stained area (“NeuN^+^ cellular area”), which provides an indication of thinning of dense neuronal layers. Lastly, an object layer was built to count individual NeuN^+^ cells. The training of this layer was performed by annotating neurons using an object size of 12μm in diameter within different brain regions. The areas included in the training present different neuronal densities and cell bodies of various morphologies and sizes to improve the robustness of the model (**Supplementary Fig. 1c & Supplementary Data 5**).

**Figure 3.**
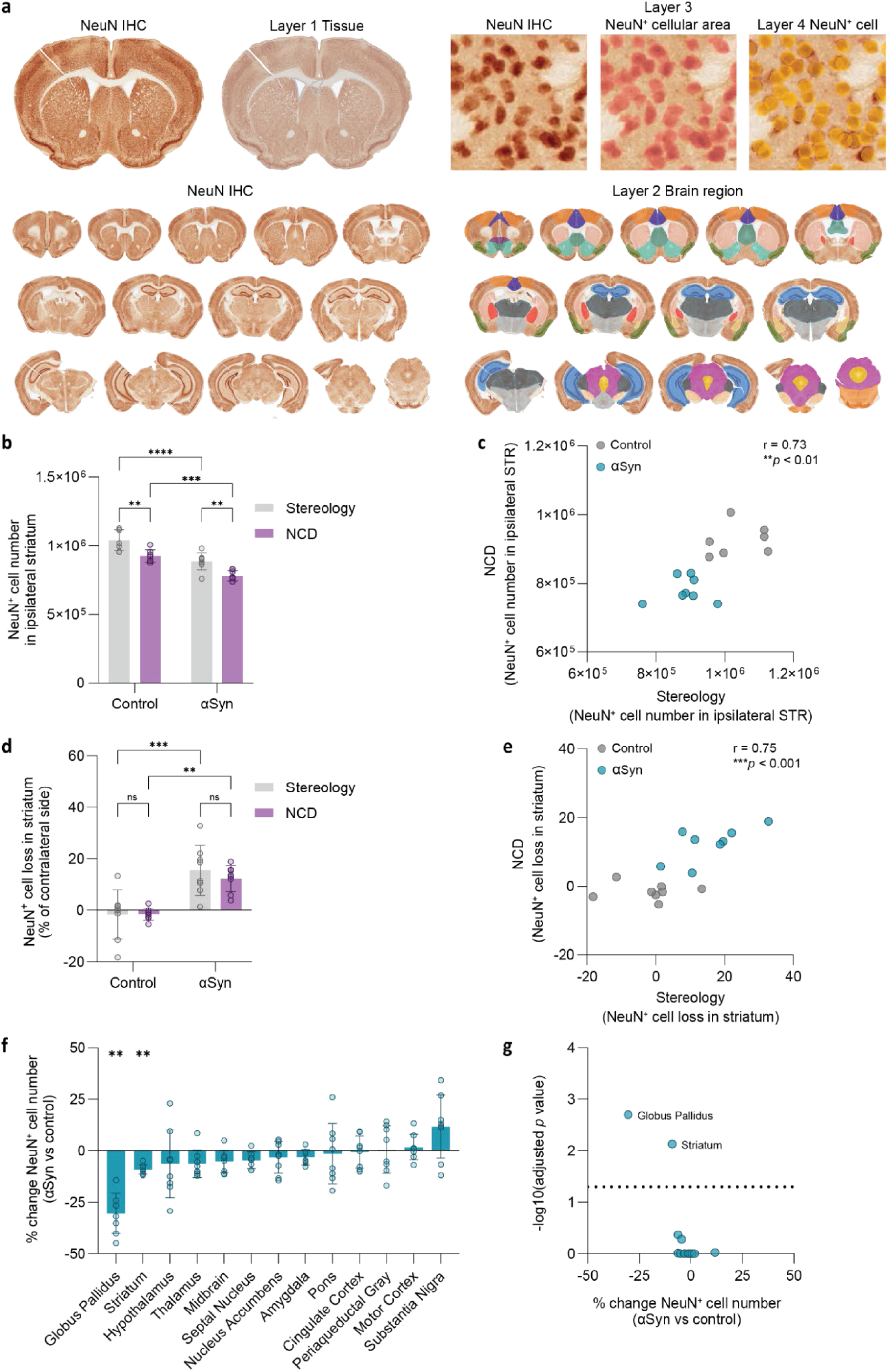
Quantification of neuronal area and neuronal cell bodies using the NCD model. Control mice were injected with PBS and αSyn mice were injected with a rAAV vector expressing αSyn in oligodendrocytes in the dorsal striatum. Analysis was performed 8 months post-injection. **a)** Examples of unannotated NeuN IHC and NCD detection per layer: “Tissue” (layer 1, gray), “Brain region” (layer 2, different color for each brain region), “NeuN^+^ cellular area” (layer 3, pale red), and “NeuN^+^ cell” (layer 4, yellow). **b)** Comparison of NeuN^+^ cell number in ipsilateral dorsal striatum measured by NCD and stereology (n = 7-8 per group, mean ± SD, two-way ANOVA test with Šídák’s correction for multiple testing, \*\**p* < 0.01, \*\*\**p* < 0.001, \*\*\*\**p* < 0.0001). **c)** Correlation between NeuN^+^ cell number in ipsilateral dorsal striatum measured by NCD and stereology (Pearson correlation r = 0.73, \*\**p* < 0.01). **d)** Comparison of dorsal striatal NeuN^+^ cell loss measured by NCD and stereology (n = 8 per group, mean ± SD, two-way ANOVA test with Šídák’s correction for multiple testing, ns, \*\**p* < 0.01, \*\*\**p* < 0.001). **e)** Correlation between NeuN^+^ cell loss in dorsal striatum measured by NCD and stereology (Pearson correlation r = 0.75, \*\*\**p* < 0.001). **f)** Brain-wide NCD analysis of NeuN^+^ cell loss with brain regions ranked according to percentage change to control group mean, each data point represents one animal (statistics performed on absolute NeuN⁺ cell numbers, n = 7-8 per group, mean ± SD, multiple Mann-Whitney tests with Holm-Šídák’s correction for multiple testing, \*\**p* < 0.01). **g)** Volcano plot of -log_10_ adjusted *p* values and percentage change of NeuN⁺ cell numbers to control group mean, each data point represents a brain region. SD (standard deviation), ns (non-significant).

A previous study showed that αSyn overexpression in oligodendrocytes leads to loss of medium spiny neurons in the dorsal striatum^25^. To evaluate the ability of the NCD model to detect neuronal loss in an animal model of MSA, we examined brain tissue from mice injected with a viral vector expressing αSyn in oligodendrocytes in the right dorsal striatum (referred to as αSyn in **Fig. 3**). Control mice were injected with PBS. Stereology was performed in the dorsal striatum of αSyn and control mice to validate the use of the NCD model. The number of ipsilateral dorsal striatal neurons in control mice was quantified with both methods and NCD found 925,217 ± 45,417 NeuN^+^ cells, while stereology found 1,040,030 ± 76,912 NeuN^+^ cells. NCD analysis of ipsilateral dorsal striatum from αSyn mice found 781,019 ± 36,685 NeuN^+^ cells while stereology found 885,836 ± 61,627 NeuN^+^ cells (**Fig. 3b**, two-way ANOVA test with Šídák’s correction for multiple testing, \*\**p* < 0.01). Despite the difference in the number of NeuN^+^ cells measured, there was significant correlation between both methods (**Fig. 3c**, Pearson’s r = 0.73, \*\**p* < 0.01). Then, we compared each method’s ability to detect a neurodegenerative lesion in the ipsilateral dorsal striatum of αSyn mice when compared to their respective contralateral side. NCD measured a 12.3 ± 5.1 % loss of NeuN^+^ cells, while stereology measured a comparable 15.5 ± 9.8 % NeuN^+^ cell loss. Control mice exhibited negligible NeuN^+^ cell loss with both methods (NCD: −1.5 ± 2.3 % NeuN^+^ cell loss; stereology: −1.7 ± 9.5% NeuN^+^ cell loss)(**Fig. 3d**, two-way ANOVA test with Šídák’s correction for multiple testing, non-significant). The quantification of NeuN^+^ cell loss significantly correlated between NCD and stereology (**Fig. 3e**, Pearson’s r = 0.75, \*\*\**p* < 0.001). These results show that despite the difference in total cell numbers detected, NCD was able to detect neurodegeneration to the same extent as stereology.

Because of the NCD model’s architecture, which contains a “Brain region” layer capable of detecting 28 different brain regions (**Supplementary Data 1**), an extensive analysis can be performed and neurons can be quantified independently within each brain region. This analysis shows that in several regions there is a reduction of NeuN^+^ cells in the αSyn-overexpressing mice. Specifically, we detect a 30.4 ± 9.7% reduction of NeuN^+^ cells in the globus pallidus and 9.1 ± 2.3% in the dorsal striatum compared to control mice (**Fig. 3f**, multiple Mann-Whitney tests with Holm-Šídák’s correction for multiple testing, \*\**p* < 0.01). Together, these results demonstrate the utility of this model, which combines sequential detection layers with an unbiased analysis approach capable of detecting brain-wide cell loss.

### Microglial cell detector

In PD and MSA, microglial alterations are observed during early stages of disease and sustained throughout the disease course. Increased microglial density and reactivity are particularly evident in regions with αSyn pathology and neurodegeneration ^26,27^. These changes affect the function and morphology of microglia, which shift from a homeostatic to a reactive state, becoming increasingly less ramified and more amoeboid in shape^28,29^.

Viral vector-mediated models of PD and MSA, in which αSyn is overexpressed in neurons or oligodendrocytes, display microglial reactivity^30,31^. To investigate neuroinflammatory changes in mouse brain we trained the microglial cell detector model, hereafter referred to as MCD, to automatically quantify Iba1^+^ microglia (**Table 2 & Fig. 4a**). Here, we further built on existing annotated layers from Stetzik and colleagues, in which a region layer, “Tissue”, and object layer, “Iba1^+^ cell”, allow for the detection of microglial cells^10^, by performing additional annotations to increase the robustness of this model. Furthermore, we incorporated a layer that identifies 28 different brain regions, creating a more efficient analysis pipeline as the regions of interest no longer need to be manually annotated (**Supplementary Data 1**). For the “Iba1^+^ cell” object layer an object size of 60μm in diameter was set. Instance segmentation was implemented in this layer to obtain accurate morphological information of individual microglial cells to assess their phenotype (**Supplementary Fig. 1d & Supplementary Data 6-7**).

**Figure 4.**
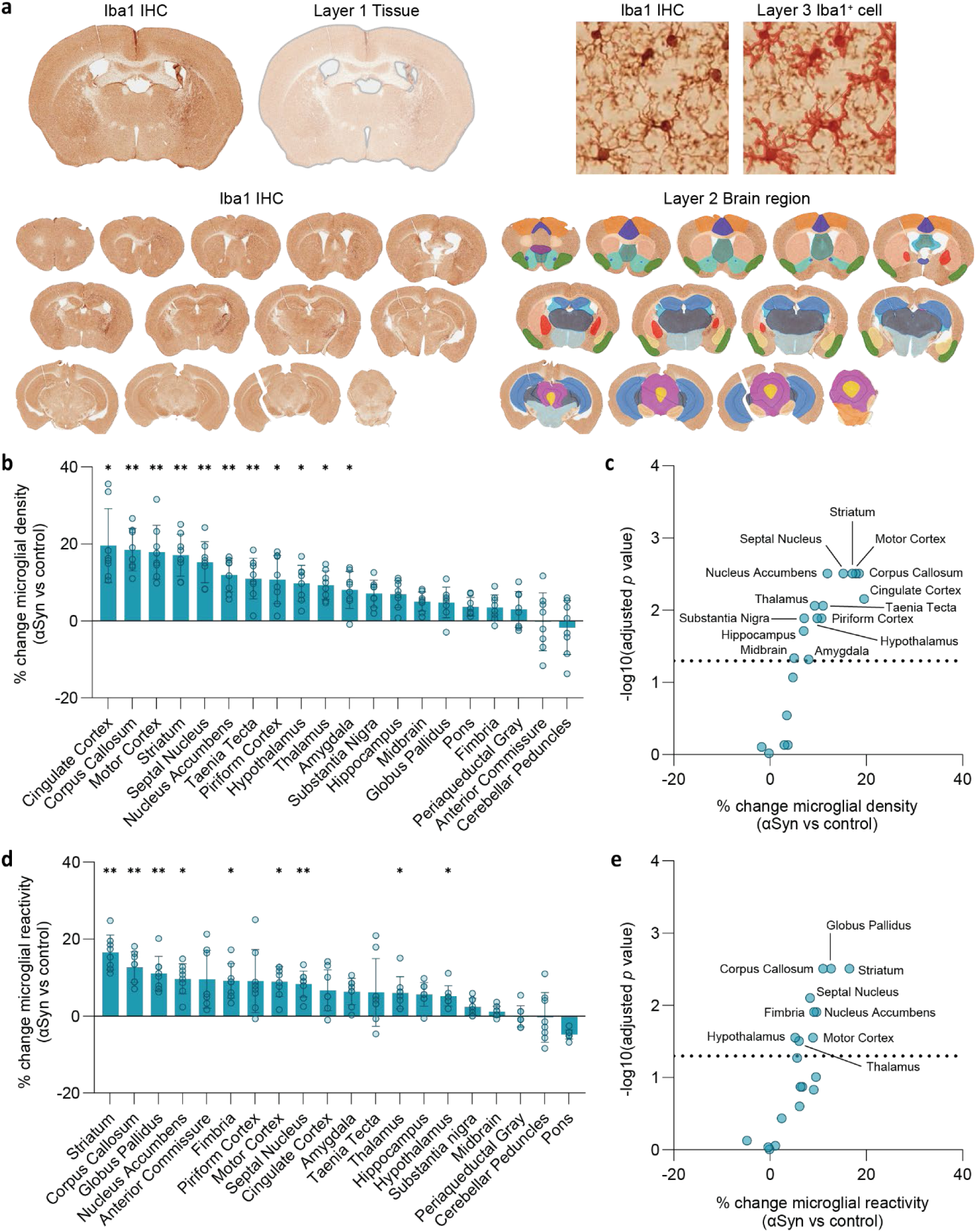
Quantification of microglial cell numbers and reactivity using the MCD model. Control mice were injected with PBS and αSyn mice were injected with a rAAV vector expressing αSyn in oligodendrocytes in the dorsal striatum. Analysis was performed 8 months post-injection. **a)** Examples of unannotated Iba1 IHC and MCD detection per layer: “Tissue” (layer 1, gray), “Brain region” (layer 2, different color for each region), and “Iba1^+^ cell” (layer 3, red). **b)** Brain-wide MCD analysis of microglial density with brain regions ranked according to percentage change to control group mean, each data point represents one animal (statistics performed on absolute Iba1⁺ cell density values, n=6-8, mean ± SD, multiple Mann-Whitney tests with Holm-Šídák correction for multiple testing, *p < 0.05, **p < 0.01). **c)** Volcano plot of -log_10_ adjusted p values and percentage change of microglial density to control group mean, each data point represents a brain region. **d)** Brain-wide MCD analysis of microglial reactivity with brain regions ranked according to percentage change to control group mean, each data point represents one animal (statistics performed on cell circularity values from Iba1⁺ cells, n = 8, mean ± SD, multiple Mann-Whitney tests with Holm-Šídák’s correction for multiple testing, *p < 0.05, **p < 0.01). **e)** Volcano plot of -log_10_ adjusted p values and percentage change of microglial reactivity (cell circularity) to control group mean, each data point represents a brain region. SD (standard deviation).

We tested the MCD model’s ability to quantify microglial cells and assess their reactive state in a viral vector-mediated MSA model expressing αSyn specifically in oligodendrocytes (referred to as αSyn in **Fig. 4**). Control mice were injected with PBS. These are the same groups analyzed previously with the NCD model. Whole brain analysis with the MCD model shows a 10 to 20 % increase in microglial density across multiple brain regions when compared to control mice. Significant changes are observed in cingulate cortex, corpus callosum, motor cortex, dorsal striatum, septal nucleus, taenia tecta, piriform cortex, hypothalamus, thalamus and amygdala (**Fig. 4b-c**, multiple Mann-Whitney tests with Holm-Šídák’s correction for multiple testing, \**p* < 0.05, \*\**p* < 0.01).

Changes in microglial density and changes in reactivity often follow distinct temporal patterns. While microglial proliferation and migration may occur independently or in advance of classical activation markers, reactivity itself can be transient or fluctuate over time, underscoring the dynamic and multifaceted nature of microglial responses in neurodegeneration^32–35^. To assess the reactive state of microglial cells the MCD model uses segmented data from the “Iba1^+^ cell” layer, which provides detailed morphometric information for all detected cells. The parameters *area* and *circumference* can then be used to derive the circularity of all microglial cells detected (**Supplementary Table 3**). Cell circularity is one of the most commonly used readouts to evaluate microglial reactivity, as it allows differentiating homeostatic ramified microglial cells (circularity values close to 0) versus amoeboid reactive ones (circularity values close to 1)^36,37^. We analyzed the morphology of microglia from control and αSyn-overexpressing mice and found that most regions with increased microglial density also exhibited significantly increased microglial reactivity (striatum, corpus callosum, motor cortex, septal nucleus, nucleus accumbens, hypothalamus and thalamus) (**Fig. 4d-e**, multiple Mann-Whitney tests with Holm-Šídák’s correction for multiple testing, \**p* < 0.05, \*\**p* < 0.01).

Overall, the MCD analysis shows there is an ongoing neuroinflammatory response in the brain of the αSyn-overexpressing mice at 8 months post-injection. While in most regions the microglial response can be detected using both density and reactivity quantifications, the incomplete overlap between both readouts in other regions could reveal a region-specific microglial phenotype. These results demonstrate the applicability of the MCD analysis pipeline to not only anatomically locate microglial cells but also to detect changes in their proliferation and phenotype.

### pSer129-αSyn detector

More than 90% of αSyn is phosphorylated at Serine 129 in the inclusions found in brains of synucleinopathy patients^11^. In PD, PSer129-αSyn is usually found in Lewy Bodies, globular-shaped dense structures in the soma of neurons, and in Lewy neurites, elongated structures in neuronal processes. MSA patients exhibit PSer129-αSyn^+^ glial cytoplasmatic inclusions in oligodendrocytes, which commonly feature a complex globular or triangular-shaped structure. This marker is also used to study the aggregation process of αSyn in animal models. When αSyn is exogenously expressed it starts filling the cytoplasm of cells, it gradually becomes phosphorylated and can be visualized as PSer129-αSyn^+^ globular shapes^38^. In seeding models, fibrillar aggregates recruit endogenous αSyn protein to form longer fibrils, which then break down, generate additional seeds, and propagate, thereby amplifying the aggregation cascade to other cells. This event typically begins in neuronal processes forming Lewy-like neurites and subsequently spreads to the soma forming globular, ring- or triangular-shaped inclusions^39–41^. In most studies, analysis of αSyn pathology is typically performed via a semi-quantitative assessment that grades the amount of inclusions per field of view or measures a percentage of PSer129-αSyn^+^ immunoreactive area, without examining quantitatively the pathological relevance of inclusion morphology^42–45^. These traditional approaches are often subject to inter-operator and inter-laboratory variability, as well as limitations due to restricted sampling.

To address these issues, we trained the pSer129-αSyn detector model, hereafter referred to as pSynD, to detect and classify brain-wide PSer129-αSyn^+^ pathology (**Table 2 & Fig. 5a**). Brain and spinal cord sections from multiple synucleinopathy mouse models and control mice processed by several researchers were included in the training to introduce variability. pSynD features a “Tissue” layer which was trained as described for previous CNN-based models. Next, this model was trained to autonomously recognize 33 different brain regions in coronal tissue sections following a similar annotation strategy as NCD and MCD models (**Supplementary Data 1**). The pSer129-αSyn^+^ inclusions are initially identified with a region layer termed “pSer129-αSyn^+^ area” that detects positively stained areas and categorizes them into neuritic inclusions (filamentous structures with neurite-like morphology) or cellular inclusions (structures with a complex globular or triangular-shaped morphology). An additional object layer, termed “Cellular inclusion”, was implemented to accurately count cellular inclusions and obtain their individual morphological metrics. Instance segmentation was used to discern overlapping cellular inclusions which can not be distinguished as two separate entities in the preceding “pSer129-αSyn^+^ area” parent layer. Cellular inclusions were annotated using an object diameter size of 7 µm and their shape was outlined using the instance segmentation tool. We did not develop an object layer for neuritic inclusions due to their intricate morphology (**Supplementary Fig. 1e & Supplementary Data 8-9**).

**Figure 5.**
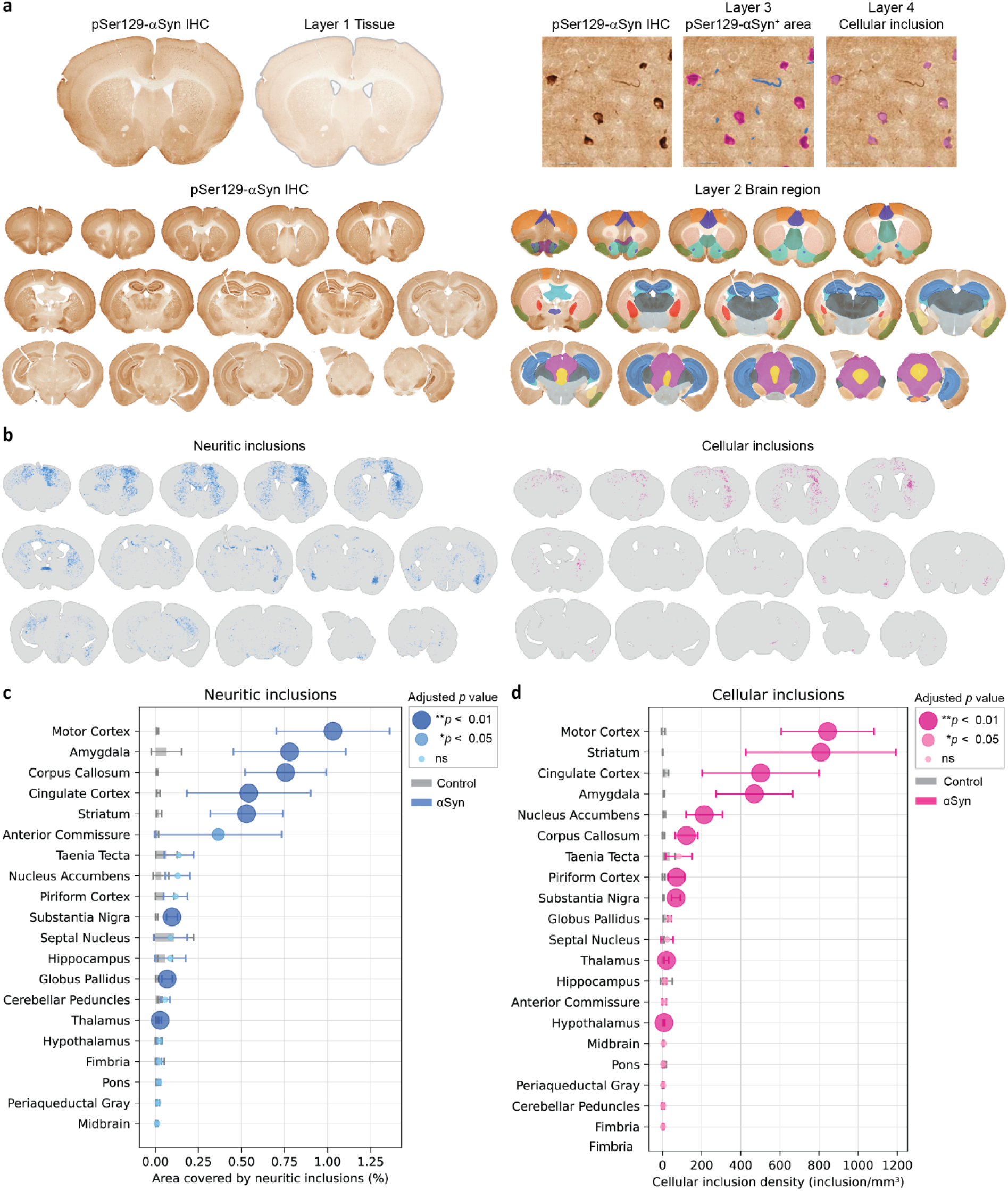
Quantification of pSer129-αSyn^+^ neuritic and cellular pathology using the pSynD model. Control mice were injected with PBS and αSyn mice were injected with PFFs in the dorsal striatum. Analysis was performed 8 months post-injection. **a)** Example of unannotated pSer129-αSyn IHC and pSynD detection per layer: “Tissue” (layer 1, gray), “Brain region” (layer 2, different color for each brain region), “pSer129-αSyn^+^ area” (layer 3, blue for neuritic inclusions and magenta for cellular inclusions) and “Cellular inclusion” (layer 4, pink). **b)** Distribution of pSer129-αSyn^+^ neuritic inclusions (left, blue) and cellular inclusions (right, magenta) in a representative PFF-injected mouse. **c)** Brain-wide pSynD analysis of pSer129-αSyn^+^ neuritic inclusions represented as percentage area covered by neuritic inclusions per brain region. Control group shown as bar and αSyn group as bubble. The size of the bubble reports the p-value difference between control and αSyn groups (n = 6-8 per group, mean ± SD, multiple Mann-Whitney tests with Holm-Šídák’s correction for multiple testing, *p < 0.05, **p < 0.01). **d)** Brain-wide pSynD analysis of pSer129-αSyn^+^ cellular inclusion density represented as number of inclusions per volume for each brain region. Control group shown as bar and αSyn group as bubble. The size of the bubble reports the p-value difference between control and αSyn groups (n = 8 per group, mean ± SD, multiple Mann-Whitney tests with Holm-Šídák’s correction for multiple testing, **p < 0.01). SD (standard deviation), ns (non-significant).

Using pSynD we examined pSer129-αSyn^+^ pathology in an αSyn seeding model. For this mouse model, PFFs are injected in the right dorsal striatum, which leads to the formation of pSer129-αSyn^+^ inclusions in the striatum and connected regions^46^. The density and location of pSer129-αSyn^+^ pathology was assessed by staining sections across the whole brain of mice injected with mouse PFFs or with PBS (referred to as αSyn and control respectively in **Fig. 5**). We observed significantly more pSer129-αSyn^+^ neuritic pathology in the αSyn mice than in the control mice in several brain regions outside the dorsal striatum, such as motor cortex, amygdala, corpus callosum, cingulate cortex, substantia nigra, globus pallidus and thalamus (**Fig. 5b-c**, multiple Mann-Whitney tests with Holm-Šídák’s correction for multiple testing, \**p* < 0.05, \*\**p* < 0.01). Interestingly, the amygdala, the taenia tecta, the piriform cortex, the septal nucleus and the hippocampus presented low but noticeable levels of endogenous pSer129-αSyn^+^ signal in control mice, observed as a speckled pattern that the pSynD model recognizes as small neurites. Despite these basal levels, pSynD was still able to detect differences in the amygdala of αSyn mice. Cellular inclusions are considered more mature forms of pathology that consolidate in the cytoplasm after aggregates relocate from neurites^39,47^. The brain regions with significantly higher density of cellular inclusions were the motor cortex and the striatum, with about 800 inclusions/mm^3^, followed by cingulate cortex, amygdala, nucleus accumbens, corpus callosum, piriform cortex, substantia nigra, thalamus and hypothalamus (**Fig. 5b, d**, multiple Mann-Whitney tests with Holm-Šídák’s correction for multiple testing, \*\**p* < 0.01).

Brain-wide analysis by the pSynD model provides a powerful yet simple tool to study spreading of PSer129-αSyn pathology. The results shown here are in line with previous studies investigating the spread of PFFs after a unilateral injection in the dorsal striatum of non-transgenic mice^48^. Besides precise anatomical mapping of PSer129-αSyn pathology, the main advantage of the pSynD model is its ability to categorize inclusions by morphology while simultaneously providing quantitative data on the absolute numbers and size of cellular inclusions, and the total area covered by neuritic inclusions. In our seeding model, regions with more neuritic inclusions do not completely correspond to regions with more cellular inclusions. This could be explained by the ratio of cellular processes versus cell bodies in specific brain regions such as the corpus callosum, a bundle of nerve fibers lacking neuronal cell bodies, in which we would expect more neuritic pathology than cellular. Alternatively, the differences could also highlight a distinct maturation status of the inclusions, regions with recent spread of pathology would contain more neuritic inclusions than cell body inclusions. We believe the pSynD model can be a valuable tool for *in vivo* studies focusing on inclusion morphology dynamics and spreading across the brain.

## Discussion

In this study we developed five CNN-based models that provide new insights in the mouse brain at cellular resolution with expert-level accuracy. Each model examines a specific neuropathological marker commonly used for histopathology of synucleinopathy, such as dopaminergic cellular and synaptic terminal pathology, neuronal changes, microglial reactivity, and pSer129-αSyn^+^ pathology. We also show how these models can be implemented to evaluate cellular changes in different anatomical regions.

The performance of the CNN-based models was evaluated by comparing their results to annotations from three experts serving as validators, using precision, sensitivity, and F1-score as validation metrics (**Supplementary Fig. 2**). Notably, there were no significant differences between the inter-validator F1-scores and the F1-scores between the models and the validators (**Supplementary Fig. 3**). This highlights the robustness and accuracy of the CNN-based models. On average, most F1-scores across all model layers exceeded 80%, indicating a high level of agreement between the models and the validators. The only exception was the “pSer129-αSyn^+^ area” layer from the pSynD model, which had a F1-score of 75% (**Supplementary Fig. 2e** & **Supplementary Fig. 4e**). This value was primarily due to a lower precision of 69%—indicating a higher rate of false positives—despite a relatively high sensitivity of 84%. This means the model overestimates the size of pSer129-αSyn^+^ structures compared to validator annotations, leading to more frequent false-positive area detections. However, there was also great disagreement among validators for this layer, with 74% precision and 74% sensitivity. This is reflected in a slightly lower inter-validator F1-score of 74%, which indicates that the 75% F1-score of pSynD versus validators is within the expected variability range for such a complex layer. Therefore, the overall expert-level performance and automation offered by the pSynD model shows it accurately quantifies pSer129-αSyn^+^ pathology and reduces inter-operator bias.

Recent studies have shown that αSyn inclusions progressively develop and evolve over time, undergoing a morphological and subcellular localization transition from neuritic filamentous structures to more compact, somatic inclusions that incorporate other components such as mitochondria and lipids. Furthermore, some compounds selectively destabilize specific αSyn inclusion types, highlighting distinct biological properties driven by their structural differences^41,49^. However, *in vivo* studies rarely interrogate the dynamic nature of αSyn aggregation or clearance, likely due to the limited availability of tools capable of distinguishing inclusion types in a quantitative and high-throughput manner. Morphological analyses of inclusions, when performed, are typically low-throughput, require sophisticated imaging (e.g., correlative light and electron microscopy, two-photon microscopy and stochastic optical reconstruction microscopy), and are more often performed for post-mortem human analysis rather than for preclinical experiments^50–52^. The pSynD analysis pipeline could tackle these limitations as it classifies and quantifies neuritic and cellular pSer129-αSyn^+^ pathology independently, making it a valuable tool for studying αSyn aggregation dynamics in longitudinal *in vivo* studies (**Fig. 5**). By following the morphological changes of neuritic and cellular inclusions over several timepoints, pSynD can assess the maturation of seeded inclusion formation and facilitate assessment of disease progression and therapeutic efficacy.

Previous studies have demonstrated the utility of AI-based analysis for human histopathology. For example, for the assessment of neuropathological markers in Alzheimer’s disease and cerebral amyloid angiopathy, kidney cancer and haematological disease diagnosis^7,53,54^. Recent studies have begun applying similar approaches for preclinical studies in mice.

However, most of these models require manual annotations or include an automated but limited brain region detection layer^8,10,48,55^. In contrast, our CNN-based models are trained to recognize a broader panel of regions, enabling detailed analysis within these defined brain regions for a more comprehensive understanding of brain-wide pathology in animal models. This approach offers significant advantages over traditional analyses, which involve pre-selecting brain regions and randomly sampling several smaller fields of view within those regions via stereology or other methods. Our CNN-based models inspect the whole area of a detected brain region, thus eliminating sampling errors. Random sampling is potentially cause of increased variability in stereological quantifications as cellular density and cell loss are not necessarily homogeneous across a brain region. In our study, NCD detected an average of 1,838,206 ± 80,073 neurons in the dorsal striatum of control C57BL/6J mice, whereas stereology detected an average of 2,026,682 ± 215,751 neurons (**Fig. 3b**). The smaller standard deviation coefficient shows that NCD provides less variable results while still yielding comparable counts to stereological values reported here and in literature^56^. Lower variability was also observed for the DCD model quantification of dopaminergic cells when compared to stereology (**Fig. 1c, e**). On average the NCD model detected fewer cells than stereology. This discrepancy could result from differences in estimation calculations performed by both methods. Alternatively, whereas stereology examines the full thickness of a section, our models detect cells or pathology on 2D-scanned images, on which no projections or serial focusing were performed. Despite this minor limitation, which could be mitigated by serial focusing (manual or z-stacking) or by using thinner tissue sections, the magnitude of decrease was consistent across control and αSyn conditions and comparable results in terms of extent of neurodegeneration were obtained (**Fig. 1e & Fig. 3d**).

Reproducibility in preclinical research is a critical factor influencing the success of clinical translation, yet it is often compromised by inconsistent analysis methods and subjective readouts across studies and research institutions^57^. Our CNN-based models directly address this challenge by providing standardized, automated analyses that produce continuous, quantitative readouts offering a more objective and reproducible alternative. For instance, the readouts from pSynD are quantitative continuous values that directly reflect the amount of pathology present in a specific brain region, rather than a semi-quantitative score based on visual inspection^42–45^. Performing statistical analyses on quantitative continuous variables offers several advantages over using discrete or categorical variables, especially in terms of sensitivity and statistical power. This consistency not only strengthens the robustness of preclinical findings but also enhances their reliability and interpretability, ultimately supporting more efficient and successful transitions to clinical trials.

Nonetheless, the vast amount of data generated when using new tools such as CNNs necessitates automated post-processing workflows. We opted to develop tailored Python code for each of the CNN-based models. These custom scripts can handle the extensive amount of information stored in each analysis file and convert it to customized metrics to gain insight in all brain regions simultaneously. This can also be performed while discriminating hemispheres in select brain regions. All Python Jupyter Notebooks are publicly available on GitHub (https://github.com/DieterPlessers/Neuro_IHC_analysis).

Even though this new methodology requires proprietary software from Aiforia®, the models developed here can be transferred to other researchers seeking to analyze the same neuropathological markers in mouse models. Furthermore, employing our CNN-based models does not require any AI-related knowledge. Despite the automation provided, we caution that inspection of brain section integrity and optimization of tissue staining methods is necessary before analysis to ensure proper performance. Nonetheless, the Aiforia® Create platform allows easily adapting the CNN-based models to new staining conditions with minimal effort, facilitating their use across institutions.

In conclusion, we have designed a high-throughput pipeline for analyzing common neuropathological markers in mouse brain, that can be shared with laboratories interested in characterizing similar readouts. The application of these novel CNN-based models for histological screening of preclinical *in vivo* models can assist researchers by reducing analysis time from weeks to minutes and providing more quantitative and unbiased readouts. We also demonstrate that these CNN-based models enable brain-wide, anatomically detailed analysis at unprecedented resolution, effectively detecting subtle neuropathological changes across multiple synucleinopathy mouse models.

## Data availability

The datasets generated and analyzed in the current study are available from the corresponding author upon reasonable request.

The underlying Python code for this study is available in GitHub as separate Jupyter Notebooks, which can be accessed via this link https://github.com/DieterPlessers/Neuro_IHC_analysis.

## Acknowledgements

The authors would like to thank J. Van Asselbergs for his technical assistance performing unbiased stereology and for validating the CNN-based models together with A. Aertgeerts, E. van Acker, E. Vonck and A. Clabout. We thank Rute Pedrosa and Darshan Kumar from Aiforia^®^ for providing technical assistance and reviewing the CNN-related methodology of this manuscript.

ABJ discloses support for this work from the FWO Flanders (doctoral fellowship 1SE8922N). EV acknowledges funding from VLAIO (Baekeland fellowship HBC.2020.2901). VB and WP disclose support for publication of this work from the FWO Flanders (G081121N, G031923N, SBO-ERA-FLAG-TRAPP MEDD S008321N) and GSKE-FMRE. The funders played no role in study design, data collection, analysis and interpretation of data, or the writing of this manuscript.

## Author contributions

ABJ and WP conceived this study. VB and WP supervised and secured funding. CVH was responsible for the viral vector production. ABJ, EV and EVA performed stereotactic injections, collected and sectioned brain tissue, performed IHC and whole-slide imaging. ABJ, EVA, WP, EV and FR contributed to the training of the CNN-based models. ABJ and DP developed Python code and performed analysis of the data and of the validation results. ABJ performed manual dopaminergic terminal quantification. ABJ drafted the manuscript and prepared the figures. All authors read and approved the final manuscript.

