## Supplementary Information for "Development of convolutional neural networks for automated brain-wide histopathological analysis in mouse models of synucleinopathies"

This file contains:

- Supplementary Figures 1-4
- Supplementary Tables 1-3
- Supplementary Data legends

**Supplementary Figure 1 Training annotations per layer for CNN-based models.** Examples of annotations are outlined in different colors for each model and layer. **a)** DCD model features three layers: layer 1 "Tissue" (green), layer 2 "Brain region" (red) and layer 3 "TH$^+$ cell" (white). **b)** DTD model features two layers: layer 1 "Tissue" (gray) and layer 2 "Brain region" (pink). **c)** NCD model features four layers: layer 1 "Tissue" (gray), layer 2 "Brain region" (different color for each brain region), layer 3 "NeuN$^+$ cellular area" (pale red), and layer 4 "NeuN$^+$ cell" (yellow). **d)** MCD model features three layers: layer 1 "Tissue" (gray), layer 2 "Brain region" (different color for each brain region), and layer 3 "Iba1$^+$ cell" (red). **e)** pSynD model features four layers: layer 1 "Tissue" (gray), layer 2 "Brain region" (different color for each brain region), layer 3 "pSer129-αSyn$^+$ area" (blue for neuritic inclusions and magenta for cellular inclusions), and layer 4 "Cellular inclusion" (pink).

**Supplementary Figure 2 Validation metrics per layer for all CNN-based models.** Precision, sensitivity and F1-score metrics calculated as average between all three validators against the performance of **a)** DCD model, **b)** DTD model, **c)** NCD model, **d)** MCD model and **e)** pSynD model.

**Supplementary Figure 3 Evaluation of model performance variability and inter-validator variability.** F1-score comparison from the annotations of three human validators against each other or the annotations of one validator against the results from **a)** DCD model, **b)** DTD model, **c)** NCD model, **d)** MCD model and **e)** pSynD model. No differences are detected between the validators and the models showing overall good performance of the models (n = 3, mean ± SD, Mann Whitney test, ns). SD (standard deviation), ns (non-significant).

**Supplementary Figure 4 Validation metrics between validators per layer for all CNN-based models*.*** Precision, sensitivity and F1-score metrics are shown in heat maps. All metrics reflect the extent to which the validations of one validator align with another, considering the first validator as the reference. Precision (TP/[TP+FP]), sensitivity (TP/ [TP +FN]), and F1-score (2 x Precision x Sensitivity/[Precision + Sensitivity]), TP (true positive), FP (false positive), FN (false negative).

**Supplementary Table 1 Post-processing exclusion criteria parameters.** Exclusion thresholds were based on the area (μm$^2$) of a subregion or object and the confidence of detection. "x" represents the area values for a subregion or object to be removed. "y" represents the confidence values for a subregion or object to be removed. If no confidence threshold is specified, exclusion is based solely on area irrespective of the confidence of detection.

**Supplementary Table 2 List of layers used by Python code to calculate readouts.**

**Supplementary Table 3 Description of readouts and calculations provided by Python code.**

Additional tables can be found in separate Supplementary Data Excel file.

**Supplementary Data 1 Layer and class hierarchy.** Describes all layers and classes included in each CNN-based model.

**Supplementary Data 2 Description of training parameters.** Describes the advanced parameters used to train the different CNN-based models. These include: i) training parameters specific to each layer, essential for training regions and/or objects, ii) image augmentation parameters, which determine the level of variability introduced to the data during training, and iii) post-training parameters, which fine-tune the resolution of the analysis results and allow reporting additional information.

**Supplementary Data 3 Training and post-training parameters for DCD model.**

**Supplementary Data 4 Training and post-training parameters for DTD model.**

**Supplementary Data 5 Training and post-training parameters for NCD model.**

**Supplementary Data 6 Training and post-training parameters for MCD model.**

**Supplementary Data 7 Gain factors for all classes in layer 2 Brain region from MCD model.**

**Supplementary Data 8 Training and post-training parameters for pSynD model.**

**Supplementary Data 9 Gain factors for all classes in layer 2 Brain region from pSynD model.**

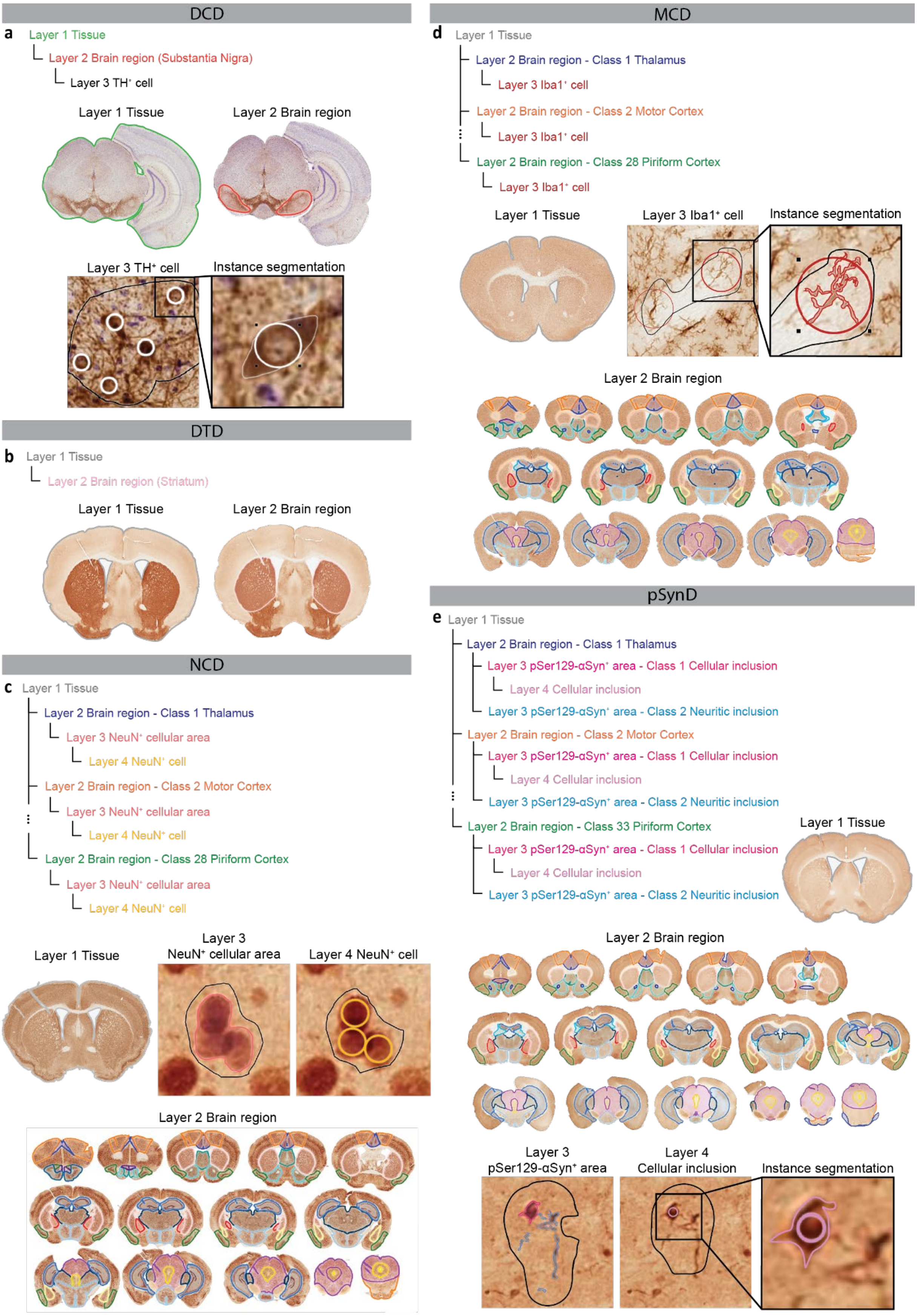


**Supplementary Figure 1 Training annotations per layer for CNN-based models.**

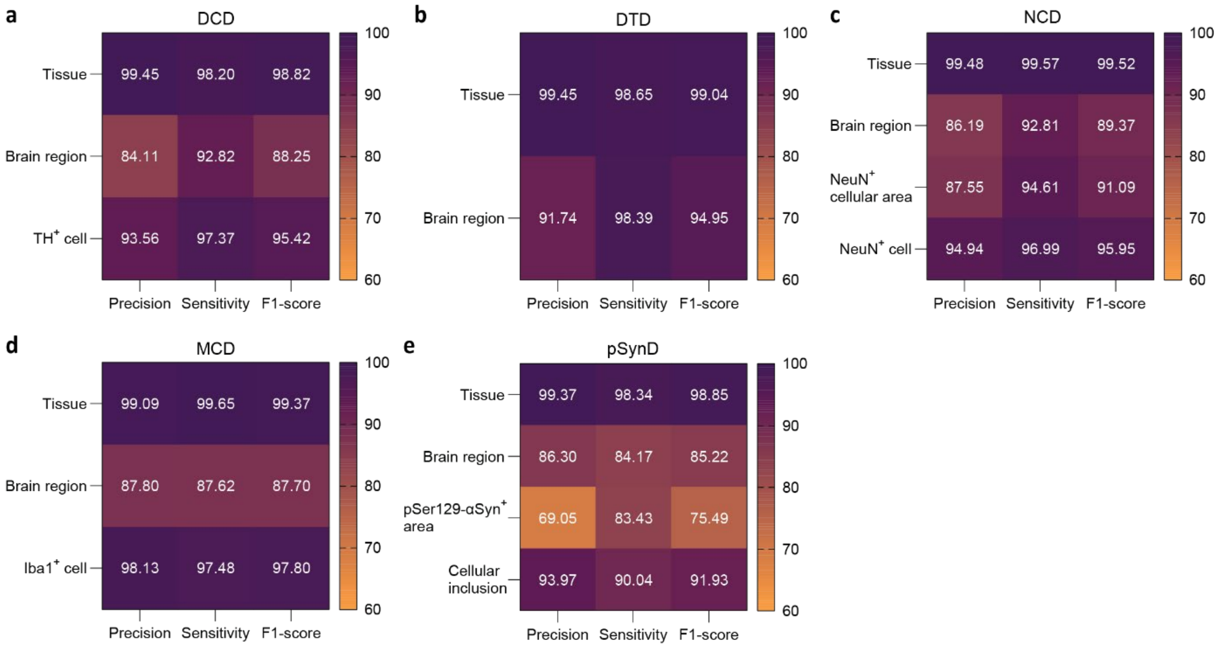


**Supplementary Figure 2 Validation metrics per layer for all CNN-based models.**

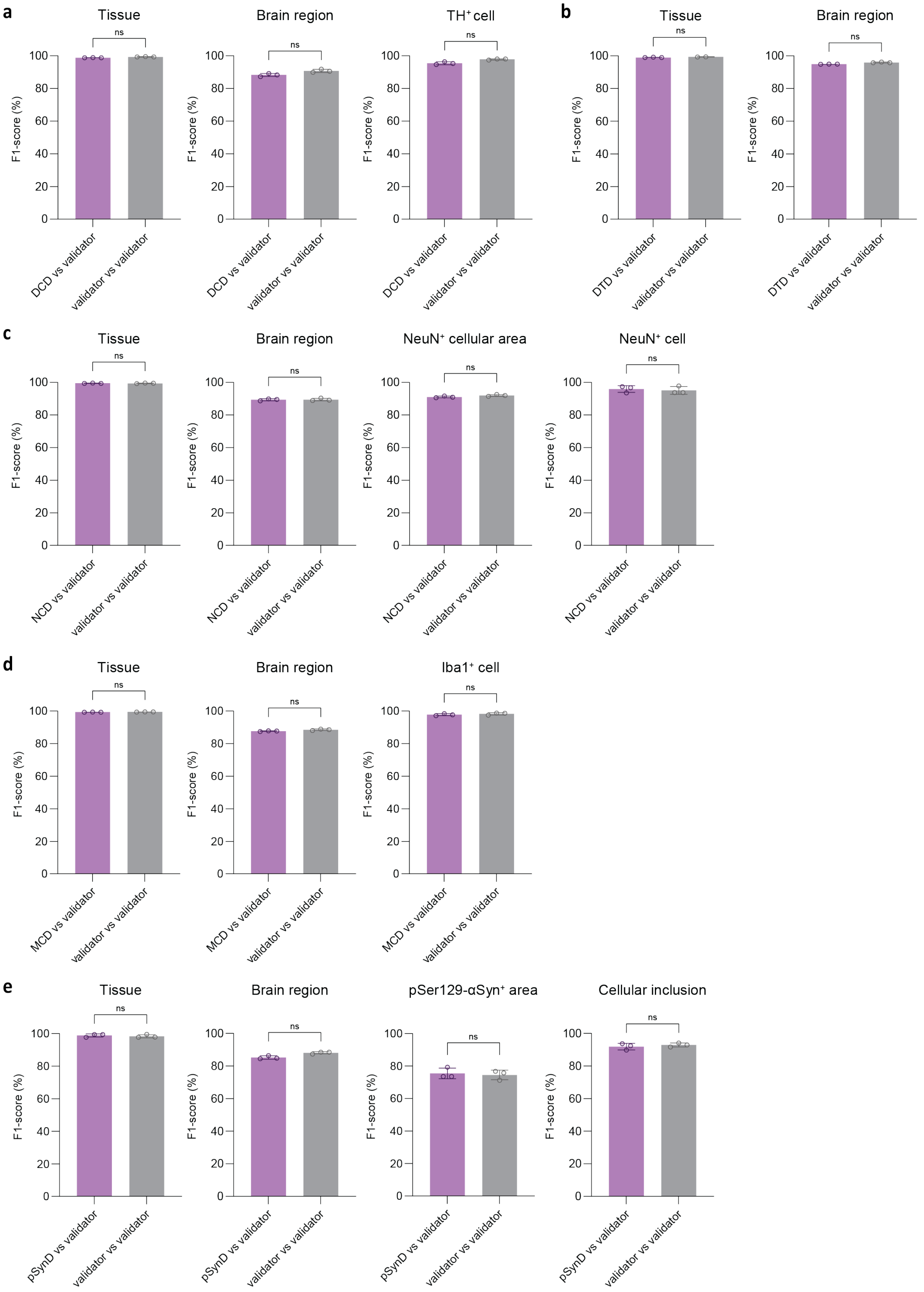


**Supplementary Figure 3 Evaluation of model performance variability and inter-validator variability.**

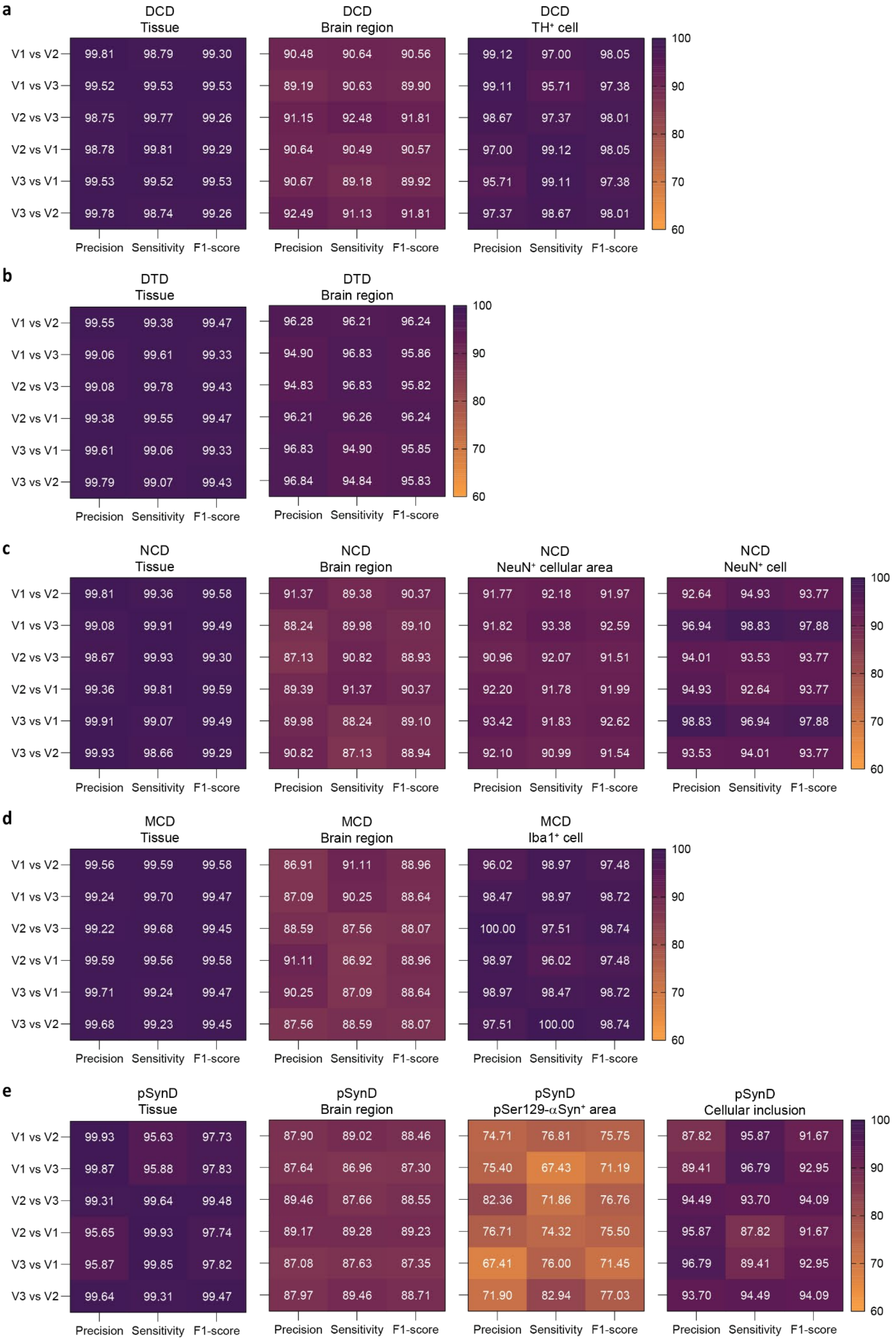


**Supplementary Figure 4 Validation metrics between validators per layer for all CNN-based models.**

**Supplementary Table 1 Post-processing exclusion criteria parameters.**

| DCD (layer 3) | DTD | NCD | MCD (layer 3) | pSynD (layers 3 & 4) |
|---|---|---|---|---|
| $x < 61$<br>$61 < x < 71$ AND $y < 80\%$<br>$71 < x < 81$ AND $y < 70\%$<br>($81 < x < 95$ OR $400 < x < 450$) AND $y < 60\%$<br>$x > 450$ | - | - | $x < 45$ | $x < 4$ (layer 3)<br>$x < 15$ (layer 4) |

**Supplementary Table 2 List of layers used by Python code to calculate readouts.**

| | DCD | DTD | NCD | MCD | pSynD |
|---|---|---|---|---|---|
| Total region area (mm$^2$) | Layer 2 | Layer 2 | Layer 2 | Layer 2 | Layer 2 |
| Weighted intensity | - | Layer 2 | - | - | - |
| Loss (%) | Layer 3 (cell number) | Layer 2 (intensity) | - | - | - |
| Average marker area (□m$^2$) | Layer 3 | - | - | Layer 3 | Layer 3 & 4 |
| Total marker area (□m$^2$) | Layer 3 | - | Layer 3 | Layer 3 | Layer 3 & 4 |
| Percentage marker positive area (%) | - | - | Layer 3 | Layer 3 | Layer 3 |
| Counts | Layer 3 | - | Layer 4 | Layer 3 | Layer 4 |
| Extrapolated counts | Layer 3 | - | Layer 4 | Layer 3 | Layer 4 |
| Counts/region area (number/mm$^2$) | Layer 3 | - | Layer 4 | Layer 3 | Layer 4 |
| Counts/region volume (number/mm$^3$) | Layer 3 | - | Layer 4 | Layer 3 | Layer 4 |
| Average area/perimeter (□m) | Layer 3 | - | - | Layer 3 | Layer 4 |
| Average circularity | Layer 3 | - | - | Layer 3 | Layer 4 |

**Supplementary Table 3 Description of readouts and calculations provided by Python code.**

| Readout | Definition | Calculation |
|---|---|---|
| Total region area (□m$^2$) | Summed area of all "$N$" subregions of the same brain region | $\sum_{i=1}^{N} Area(brain\ subregion_i)$ |
| Weighted intensity | Average weighted intensity of all "$N$" subregions of the same brain region (weights based on subregion area) | $\frac{\sum_{i=1}^{N} Intensity(brain\ subregion_i) \times Area(brain\ subregion_i)}{\sum_{i=1}^{N} Area(brain\ subregion_i)}$ |
| Loss (%) | Percentage comparing contralateral versus ipsilateral values | $100\%\ x\ (1 - \frac{ipsilateral}{contralateral})$ |
| Average marker area (□m$^2$) | Average area of all "$n$" marker subregions OR all "$M$" object areas within a brain region | $\frac{\sum_{i=1}^{n} Area(marker\ subregion_i)}{n}$ <br> $\frac{\sum_{i=1}^{M} Area(object_i)}{M}$ |
| Total marker area (□m$^2$) | Summed area of all "$n$" marker subregions OR all "$M$" object areas within a brain region | $\sum_{i=1}^{n} Area(marker\ subregion_i)$ <br> $\sum_{i=1}^{M} Area(object_i)$ |
| Percentage marker positive area (%) | Percentage of marker positive area within a brain region | $\frac{Total\ marker\ area\ (\mu m^2)}{Total\ region\ area\ (\mu m^2)}\ x\ 100\%$ |
| Counts | Number of "$M$" objects detected within a brain region | $\mathrm{M} = \text{counts of objects}$ |
| Extrapolated counts | Number of total objects estimated within a brain region | $\text{Serial section spacing interval} \times \mathrm{M}$ |
| Counts/region area (number/mm$^2$) | Object density within a brain region | $\frac{M}{Total\ region\ area\ (\mu m^2)}\ x\ 10^6$ |
| Counts/region volume (number/mm$^3$) | Object density within a brain region volume | $\frac{M}{Total\ region\ area(\mu m^2) \times Section\ thickness(\mu m)}\ x\ 10^9$ |
| Average area/perimeter (□m) | Parameter to assess morphology of all "$M$" objects within a brain region | $\frac{\sum_{i=1}^{M} \frac{Area(object_i)}{Circumference(object_i)}}{M}$ |
| Average circularity | Parameter to assess roundness of all "$M$" objects within a brain region | $\frac{4 \times \pi \times \sum_{i=1}^{M} \frac{Area(object_i)}{Circumference(object_i)^2}}{M}$ |
